# The predator activity landscape predicts the anti-predator behavior and distribution of prey in a tundra community

**DOI:** 10.1101/2020.10.16.342725

**Authors:** Jeanne Clermont, Alexis Grenier-Potvin, Éliane Duchesne, Charline Couchoux, Frédéric Dulude-de Broin, Andréanne Beardsell, Joël Bêty, Dominique Berteaux

## Abstract

Predation shapes communities through consumptive and non-consumptive effects, where in the latter prey respond to perceived predation risk through risk management strategies occurring at different spatial and temporal scales. The landscape of fear concept is useful to better understand how predation risk affects prey behavioral decisions and distribution, and more generally the spatial dimension of predator-prey relationships. We assessed the effects of the predation risk landscape in a terrestrial Arctic community, where arctic fox is the main predator of ground-nesting bird species. Using high frequency GPS data, we developed a predator activity landscape resulting from fox space use patterns, and validated with an artificial prey experiment that it generated a predation risk landscape. We then investigated the effects of the fox activity landscape on multiple prey, by assessing the anti-predator behavior of a primary prey (snow goose) and the nest distribution of several incidental prey. Areas highly used by foxes were associated with a stronger level of nest defense by snow geese. We further found a lower probability of occurrence of incidental prey nests in areas highly used by foxes, but only for species nesting in habitats easily accessible to foxes. Species nesting in refuges consisting of micro-habitats limiting fox accessibility, like islets, did not respond to the fox activity landscape. Consistent with the scale of the fox activity landscape, this result reflected the capacity of refuges to allow bird nesting without regard to predation risk in the surrounding area. We demonstrated the value of using predator space use patterns to infer spatial variation in predation risk and better understand its effects on prey in landscape of fear studies. We also exposed the diversity of prey risk management strategies, hence refining our understanding of the mechanisms driving species distribution and community structure.

## Introduction

Predation plays a central role in ecological and evolutionary processes (Menge and Sutherland 1976, Ford et al. 2014). It shapes communities through both direct killing of prey (consumptive effects) and triggering of costly anti-predator responses (non-consumptive effects) (Lima and Dill 1990, Cresswell 2008, Laundré et al. 2010). Non-consumptive effects of predation can be major drivers of food web structure and dynamics (Cresswell 2008, Teckentrup et al. 2018).

Prey respond to predation risk with various risk management strategies at broad, intermediate and fine spatial and temporal scales (Lima and Dill 1990, Guiden et al. 2019). At broad scales (kilometers, days), predation risk associated to different areas may influence prey’s choice of home range, such as the breeding home range of migrant birds (Lima 2009, Morosinotto et al. 2010). At intermediate scales (hectares, hours), variation in predation risk within the home range of a prey may affect its space use. For example, many bird species maximise their reproduction by nesting where predation risk is the lowest, either where the regional abundance of main predators is low (Forstmeier and Weiss 2004) or in habitats providing refuge against predation (Anderson et al. 2015). Such refuges can offer complete or partial protection. At fine scales (meters, minutes), when predator encounter is imminent, prey use anti-predator behavior such as escape behavior (Ydenberg and Dill 1986). In many species, parents (such as incubating birds) also provide offspring defense (Montgomerie and Weatherhead 1988, Lima 2009).

Prey risk management strategies also depend on predator and prey encounter rates (Gaynor et al. 2019). Indeed, space use patterns of predators and their primary prey (which are often the most abundant and profitable prey, Stephens and Krebs 1986) tend to correlate (Fortin et al. 2005, Arias-Del Razo et al. 2012). Thus, primary prey species can hardly avoid predation risk by shifting their home ranges because predators actively search for them. Prey can rather adopt fine scale risk management strategies such as defense behaviors or increased vigilance when using risky areas (Laundré et al. 2001). On the other hand, incidental prey species, which are consumed when encountered but are not actively searched, may manage risk of predation by avoiding areas highly used by predators (Forstmeier and Weiss 2004, Avgar et al. 2015).

The landscape of fear concept offers a useful framework to understand how predation risk affects prey behavior (Laundré et al. 2010, Gaynor et al. 2019). Laundré et al. (2010) defined the landscape of fear as the spatial variation in prey perception of predation risk. Gaynor et al. (2019) then framed the landscape of fear as part of a series of interdependent landscapes. First, the physical landscape represents habitat features that interact with the biology (hunting mode, body size, etc.) of predators and prey to determine their distributions and interactions. These interactions then modulate the predation risk landscape and, accordingly, the landscape of fear. Finally, the landscape of fear determines the responses of prey to predation risk, which ultimately shape spatiotemporal variations in prey distribution and anti-predator behavior. Many studies have used proxies of predation risk, such as habitat features (Dupuch et al. 2014), or proxies of perceived predation risk, such as prey behavior (Willems and Hill 2009). Proxies are useful but they can also lead to circular reasoning (Gaynor et al. 2019).

In active hunting predators, space use of active individuals, which can be measured at a fine scale through GPS tracking, should closely approximate the landscape of predation risk since they continuously prowl in search of prey (Schmitz et al. 2004). Some landscape of fear studies measured predator movements to explain prey behavior while considering local density or space use of predators, but with only a limited number of locations (Thaker et al. 2011, Kohl et al. 2018). For very active predators, a detailed assessment of movements is required to infer the predation risk landscape (Poulin et al. 2020). Fortunately, improved data acquisition (Wilmers et al. 2015) and modelling techniques (e.g., hidden Markov models, Patterson et al. 2017) now allow to assess the behavior and active periods of predators from their fine scale movements. However, the validity of using fine scale predator space use patterns as a surrogate to the predation risk landscape should be demonstrated rather than assumed.

To better understand the effects of the landscape of fear on natural communities, we need to simultaneously evaluate how predators generate the distribution of predation risk and how prey respond to this distribution, ideally including all important predators and prey of the system (Gaynor et al. 2019). Arctic terrestrial food webs are good models to study vertebrate predator-prey interactions because they are relatively simple. One example is the tundra community of Bylot Island (Nunavut, Canada), where the arctic fox (*Vulpes lagopus*) is the main terrestrial predator. This canid is an active hunting predator that travels extensive daily distances within its territory (Poulin et al. 2020). On Bylot, it feeds primarily on lemmings (*Lemmus trimucronatus* and *Dicrostonyx groenlandicus*) which show important annual density fluctuations (Gruyer et al. 2008). During summer, foxes also collect eggs of the colonial nesting greater snow goose (*Anser caerulescens antlanticus*) for immediate consumption and for storage (Bêty et al. 2001). As such, foxes select in summer productive lemming habitats and patches of high snow goose nest density (Grenier-Potvin et al. 2020). Because snow geese cannot avoid areas highly used by foxes, they actively defend their nests when closely approached by a fox (Bêty et al. 2002). This defense strategy is effective as long as geese remain close to their nest during incubation (Reed et al. 1995). Foxes also opportunistically prey upon nests of other ground nesting birds and are their main nest predator (McKinnon and Bêty 2009, Gauthier et al. 2011). These incidental prey mainly nest in mesic tundra, but some nest in micro-habitats that constrain fox movements and thus offer protection (Lecomte et al. 2008). For example, islets of just a few spare meters located in ponds may serve as refuges (Gauthier et al. 2015).

We assessed the effects of the predation risk landscape in the tundra community of Bylot Island. We first defined and assessed empirically the *predator activity landscape*, that is the utilization distribution of active foxes, using high frequency GPS data coupled with hidden Markov models. We then experimentally tested if this predator activity landscape predicted (P1) the probability of consumption of artificial prey, thus reflecting the predation risk landscape. Then, we investigated the effect of the fox activity landscape on risk management strategy and nest distribution of the bird community. We assessed the nest defense behavior of a primary prey (snow geese), predicting (P2) that nest defense would be stronger in areas most used by foxes, where predation risk is higher. We also assessed the effect of the predator activity landscape on the nest distribution of incidental prey (P3), composed of bird species from different guilds. For species nesting in common habitats easily accessed by foxes, we predicted (P3a) that the probability of nest occurrence should be lowest in areas most used by foxes. Given the limited spatial resolution of the fox activity landscape, we predicted that for species nesting in small refuges such as islets, the probability of nest occurrence should be independent of the fox activity landscape (P3b), as the location of refuges used for nesting should be independent of the predation risk in the surrounding landscape.

## Methods

### Study system

We worked during summer 2019 in the southwest plain of Bylot Island (72°53’ N, 79°54’ W), in Sirmilik National Park of Canada, Nunavut (Appendix S1: Fig. S1). The ecosystem is characterized primarily by mesic tundra and polygonal wetlands (Grenier-Potvin et al. 2020). In this system, arctic fox pairs have virtually no predators and are territorial. All studied individuals had their territory in a snow goose colony composed of > 20,000 nesting pairs distributed over 70 km^2^(Bêty et al. 2001, Bêty et al. 2002). Lemming density was high enough to allow reproduction of 5 of the 6 monitored fox pairs.

### Fox captures and movement tracking

During May and June 2019, 13 foxes were captured using Softcatch #1 padded leghold traps (Oneida Victor Inc. Ltd., Cleveland, OH, USA). These foxes represented 6 neighboring territorial pairs and one additional individual, whose small home range overlapped two territories (Fig. 1). Each fox was marked with colored ear tags allowing identification at a distance, and was fitted with a GPS collar (95 g, 2.6–3.3% of body mass; Radio Tag-14, Milsar, Poland) equipped with rechargeable batteries, a solar panel, and UHF transmission allowing remote data download. We used a GPS fix interval of 4 min and average GPS location error was 11 m (Poulin et al. 2020). The 6 fox territories represent our study area. The general contour of the study area was drawn using the concave hull of fox GPS data (QGIS version 3.8.3, QGIS Development Team 2019), excluding a few extra-territorial trips (Fig. 1). For each individual, we used locations from 10 days at the end of June. Datasets were synchronized (± 2 days depending on capture day and the timing of missing data; the 2 days following capture were excluded) and matched laying and incubation of birds. Daily observations and automated cameras at fox dens confirmed that we tracked all foxes foraging in the study area.

**Fig. 1.**
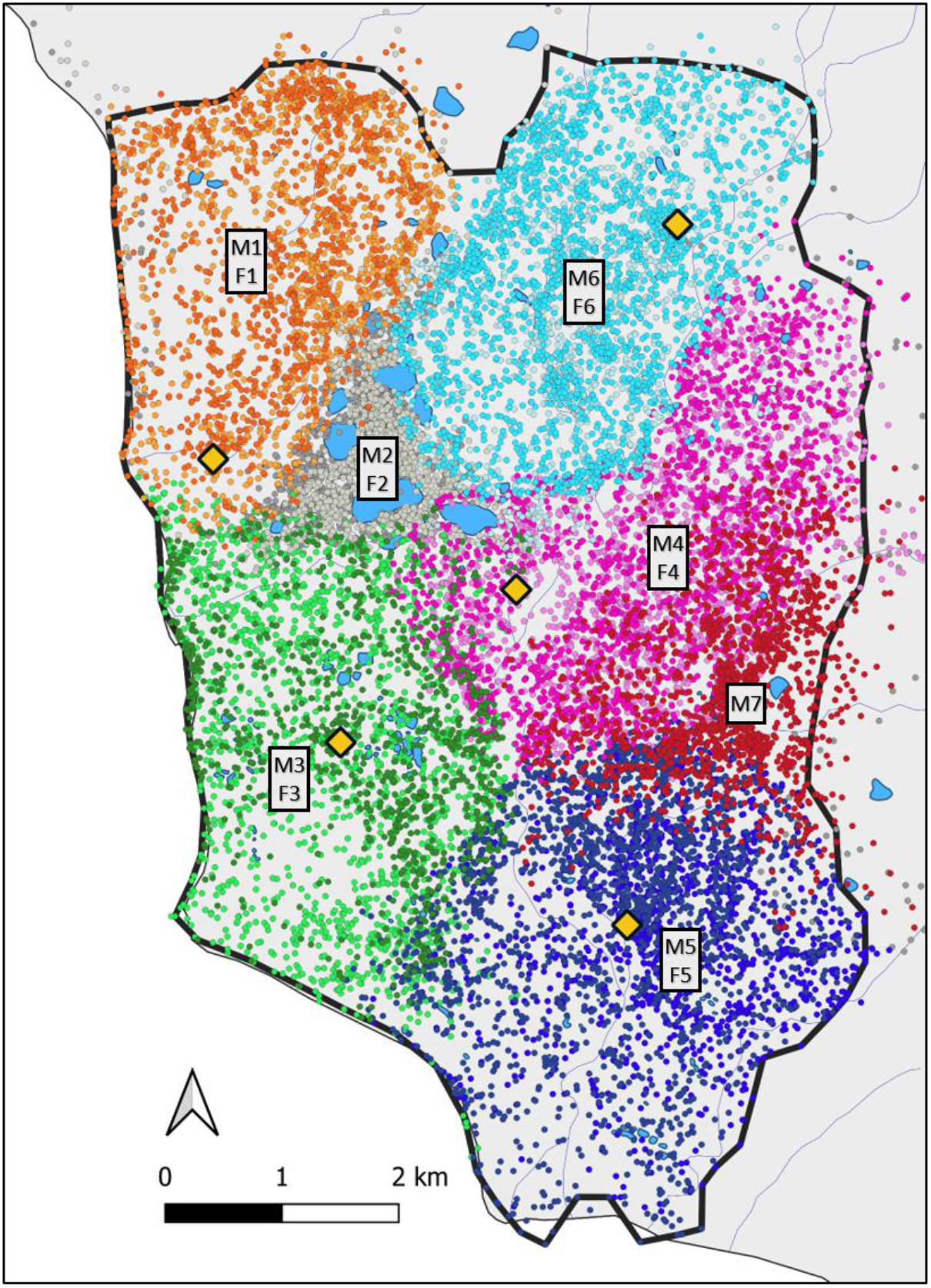
Study area on Bylot Island (Nunavut, Canada) featuring GPS locations of 13 arctic foxes tracked during 10 days at the end of June 2019. Foxes occupied 6 territories (M7 had a small home range overlapping two territories) and pair members have related colors, as detailed in Appendix S1: Table S1. GPS locations were collected at 4-min fix intervals and the 20,961 data points shown are those classified in the active state by a hidden Markov model. Yellow diamonds locate the 5 reproductive dens (M2, F2 and M7 did not reproduce). The thick black line is the contour of the study area. Lakes and large ponds are in blue. See Appendix S1: Fig. S1 for the geographical context of the study area.

Capture techniques and immobilization procedures were approved by the UQAR Animal Care Committee (CPA-64-16-169 R3) and field research was approved by the Joint Park Management Committee of Sirmilik National Park of Canada (SIR-2018-28021).

### Artificial prey experiment

We conducted an artificial prey experiment using 8-g pieces of dried beef liver (measuring ca. 0.5 × 2 × 2 cm; Benny Bullys Sales Inc., ON, Canada; hereafter, baits) to assess predation risk. On 4 July, we placed in each of the 6 fox territories 6–8 lines of ca. 10 baits each (total of 428 baits distributed in 44 curved lines each measuring 293 ± 77 m (mean ± SD), see baits locations in Appendix S1: Fig. S2A). Each bait line was located in a homogeneous habitat patch and bait lines were distributed equally between wetland polygons and mesic tundra patches, at least 300 m from the study area limits to avoid potential border effects. Distance between adjacent baits within bait lines was of 79 ± 7 m and distance between adjacent lines was 297 ± 118 m. Baits were covered with moss or lichen to exclude predation by avian predators (as done for artificial nests, Léandri-Breton and Bêty 2020) and were visited after 4 days to assess their removal by foxes. A piece of orange flag placed under each bait became visible when a bait had been removed, thus facilitating assessment of predation events. After the experiment, camera traps were placed during 5 ± 2 days at 6 locations (in 3 fox territories) to monitor the fate of baits, which were replaced if consumed. Thirteen baits were taken, always by foxes, thus confirming this species as the only bait consumer.

### Snow goose nest defense behavior

The threat posed by predators is often much higher for young than for adults (Rosenbaum 2018). Adults from species with large body size are almost immune to predation and therefore defend their offspring rather than flee (Rosenbaum 2018). Flushing distance from an approaching human is often used to assess a prey anti-predator strategy (Blumstein 2003) and represents a good proxy for nest defence intensity. On Bylot, foxes essentially pose a threat to snow goose eggs and chicks (Bêty et al. 2001), so we measured the flushing distances of 458 incubating females as an indicator of their level of nest defense (see nest locations in Appendix S1: Fig. S2B). A small flushing distance (the observer is close to the nest when the female leaves) indicates a high level of nest defense (Clermont et al. 2019). An observer approached a focal nest by walking silently at a slow and constant pace, in a straight line, and measured flushing distance with a telemeter or handheld GPS. To limit potential effects of incubation stage on goose nest defense (Clermont et al. 2019), we performed 85% of flushing distance measures within 5 days from June 14 to June 18 (we did remaining measures in the following days), which corresponds to the first half of the incubation period. We also assessed clutch size which generally influences nest defense intensity (Montgomerie and Weatherhead 1988), and we measured the starting distance of the approach which affects flushing distance (Blumstein 2003). Focal nests were located at least 300 m from the study area limits to avoid potential border effects.

### Nest distribution of incidental prey

During the incubation period, we conducted thorough searches of bird nests other than snow geese (i.e., incidental prey). In June, we walked repeatedly throughout the study area to detect signs of reproductive birds (calling, distraction displays, bird flushing at close distance). This was done through transect surveys conducted in mesic tundra, and intensive nest searches performed in wetland patches, stony riverbanks and slopes, which are all easily accessible to foxes. We also inspected enclaves, mostly islets in ponds, with a few peninsulas located in complex wetlands, where nests are less accessible to foxes as they are surrounded by water. We georeferenced 377 islets in the study area (Appendix S1: Fig. S3).

We found 109 nests from 13 species in the study area (see nest locations in Appendix S1: Fig. S2C). A total of 44 nests from 10 species were located in areas easily accessible to foxes: common-ringed plover (*Charadrius hiaticula*, n = 3), american golden plover (*Pluvialis dominica*, n = 9), white-rumped sandpiper (*Calidris fuscicollis*, n = 2), arctic tern (*Sterna paradisaea*, n = 2), rough-legged hawk (*Buteo lagopus*, n = 1), lapland longspur (*Calcarius lapponicus*, n = 16), parasitic jaeger (*Stercorarius parasiticus*, n = 1), long-tailed jaeger (*Stercorarius longicaudus*, n = 6), long-tailed duck (*Clangula hyemalis*, n = 1) and king eider (*Somateria spectabilis*, n = 3). A total of 65 nests from 3 species were located in refuges: cackling goose (*Branta hutchinsii*, n = 38), glaucous gull (*Larus hyperboreus*, n = 11) and red-throated loon (*Gavia stellate*, n = 16).

### Predator activity landscape

We defined the predator activity landscape as the utilization distribution (see below) of all foxes in the active state within the study area. For opportunist active hunting predators like arctic foxes, all travelling phases can be associated with hunting, therefore we used a hidden Markov model (HMM) to assign GPS locations to an active or resting state (R package moveHMM, Michelot et al. 2016). HMM decomposes GPS tracks into sequences associated to different behavioral states, which differ from one another in their step lengths and turning angles (Langrock et al. 2012). The active state is characterized by long step lengths and small turning angles, and the resting state with short step lengths and large turning angles. The HMM included time of the day as a covariate to reflect the circadian rhythm of foxes (Grenier-Potvin et al. 2020). Models using a Weibull distribution for step lengths and a wrapped Cauchy distribution for turning angles yielded the most parsimonious model (HMM construction and model selection is detailed in Grenier-Potvin et al. 2020).

Then, we used Kernel Density Estimation (QGIS Heatmap plugin) to map the fox utilization distribution (UD) using only active locations. UDs quantify the intensity of space use (from low to high probability density of GPS locations) by tracked animals and thus identify areas where animals are most likely to be found (Fortin et al. 2005, Thaker et al. 2011). We used 10-m^2^pixels to map UD scores, and a fixed UD smoothing parameter (called radius in QGIS, which is equivalent to the kernel bandwidth) to specify the distance at which GPS locations influence UD scores. As the choice of the UD smoothing parameter can affect prediction tests, we performed a sensitivity analysis. We ran statistical models (presented in the following section) for 5 UD smoothing parameters ranging from 200 to 400 m (50-m increments). As foxes in their active state traveled 232 ± 145 m (mean ± SD, see Results) between GPS fixes obtained at 4-min intervals, the chosen range of smoothing parameters yielded fine resolution activity landscapes that reflected the scale of our data. Using smaller parameter values would have underestimated the use of areas located between GPS locations, whereas using larger parameter values would have overestimated the use of areas located on each side of the fox track. UD scores were standardized from 0 to 1 in each of the 5 UDs.

### Statistical models

We tested the effect of the fox activity landscape on the probability of predation of baits (P1), snow goose nest defense behavior (P2) and the nest distribution of fox incidental prey (P3). A first step consisted in extracting the fox UD score at all locations used in the models, that is locations of baits, nests of tested snow geese, and nests and available nesting locations of incidental birds (see below).

#### 1) Probability of predation of baits

We used a generalized linear mixed model (R package lme4, Bates et al. 2015) with a logit-link function and a binomial distribution to test the effect of fox UD score on the probability of predation of baits (0 = not predated, 1 = predated), with the ID of the bait line nested in the ID of the fox territory as random effects. We fitted one model for each of the 5 UDs defined with different smoothing parameters.

#### 2) Snow goose nest defense behavior

We used a linear mixed model to test the effect of fox UD score on goose flushing distance. Goose flushing distance was square-root transformed to respect the assumption of normality and homoscedasticity in models’ residuals. The other fixed effects included in the models were clutch size, starting distance of the approach, and date of observation. All covariates were centered and standardized to facilitate interpretation of model estimates (Schielzeth 2010). We included as random effects the ID of the fox territory and the ID of the observer performing the approach. We fitted one model for each of the 5 UDs.

#### 3) Nest distribution of incidental prey

We used conditional logistic regressions with a use-available design (function clogit in R package survival, Therneau et al. 2020) to test the effect of fox UD score on the distribution of bird nests of fox incidental prey species. We analyzed separately species nesting in habitats easily accessible to foxes (first set of models, P3a) and species nesting in refuges (second set of models, P3b).

In the first set of models, we compared fox UD scores at bird nests (used locations) to fox UD scores at random sites (available locations). We considered as available locations potential nesting sites located in the study area, excluding water bodies. Each bird nest location was paired to 50 random locations drawn from an area surrounding the nest (hereafter, the *nest area*). As tundra nesting birds have various natural histories, including nesting habitat and social system, they likely select nesting sites at different spatial scales which are unknown. Hence, we could not justify *a priori* a single radius for the nest area. We therefore repeated analyses after forcing random locations within 5 radii varying from 1000 m to 3000 m (increments of 500 m), thus fitting 25 models (5 UDs × 5 nest area radii).

In the second set of models, we again compared fox UD scores at bird nests (used locations) to fox UD scores at available sites. However, we used as available locations potential nesting sites located in the study area and surrounded by water, drawing from our 377 georeferenced islets. Each bird nest location was paired to 50 islets chosen randomly from the area surrounding the nest. Less than 50 islets were sometimes available, so we assessed whether this affected results (Appendix S2). Since fox UD scores are smoothed values obtained from GPS locations collected at a 4-min fix interval, they reflect fox utilization of the surrounding area rather than micro-habitat use, and the UD score of an islet could be > 0 even if no fox visited the islet. As for the first set of models, we fitted 25 models (5 UDs × 5 nest area radii).

All analyses were conducted in R (version 3.6.1, R Development Core Team 2019). We validated the assumptions of normality, homoscedasticity, non-collinearity among fixed effects, and independence of residuals for all models. Values are expressed as mean ± SD.

## Results

### Fox activity landscape

A total of 45,140 fixes were acquired for 13 foxes tracked for 10 days (Fig. 1). The active behavioral state was assigned to 46 ± 9% of locations per individual (range 31–60%, Appendix S1: Table S1) for a total of 20,961 GPS locations. Average step length and turning angle were 232 ± 145 m and 55° for active locations, and 9 ± 9 m and 116° for resting locations (see Grenier-Potvin et al. 2020 for detailed HMM results). The representation of the fox activity landscape (Fig. 2, UD smoothing parameter = 300 m) shows obvious heterogeneity in the intensity of space use by foxes. This heterogeneity decreased, but overall patterns remained as smoothing parameters varied from 200 m to 400 m (Fig. 2, Appendix S1: Fig. S4). The predator activity landscape identified areas intensively used by some foxes, such as the small central territory where individuals M2 and F2 lived in a restricted area, and sections M7 shared with M4, F4, M5 and F5 (Figs. 1, 2).

**Fig. 2.**
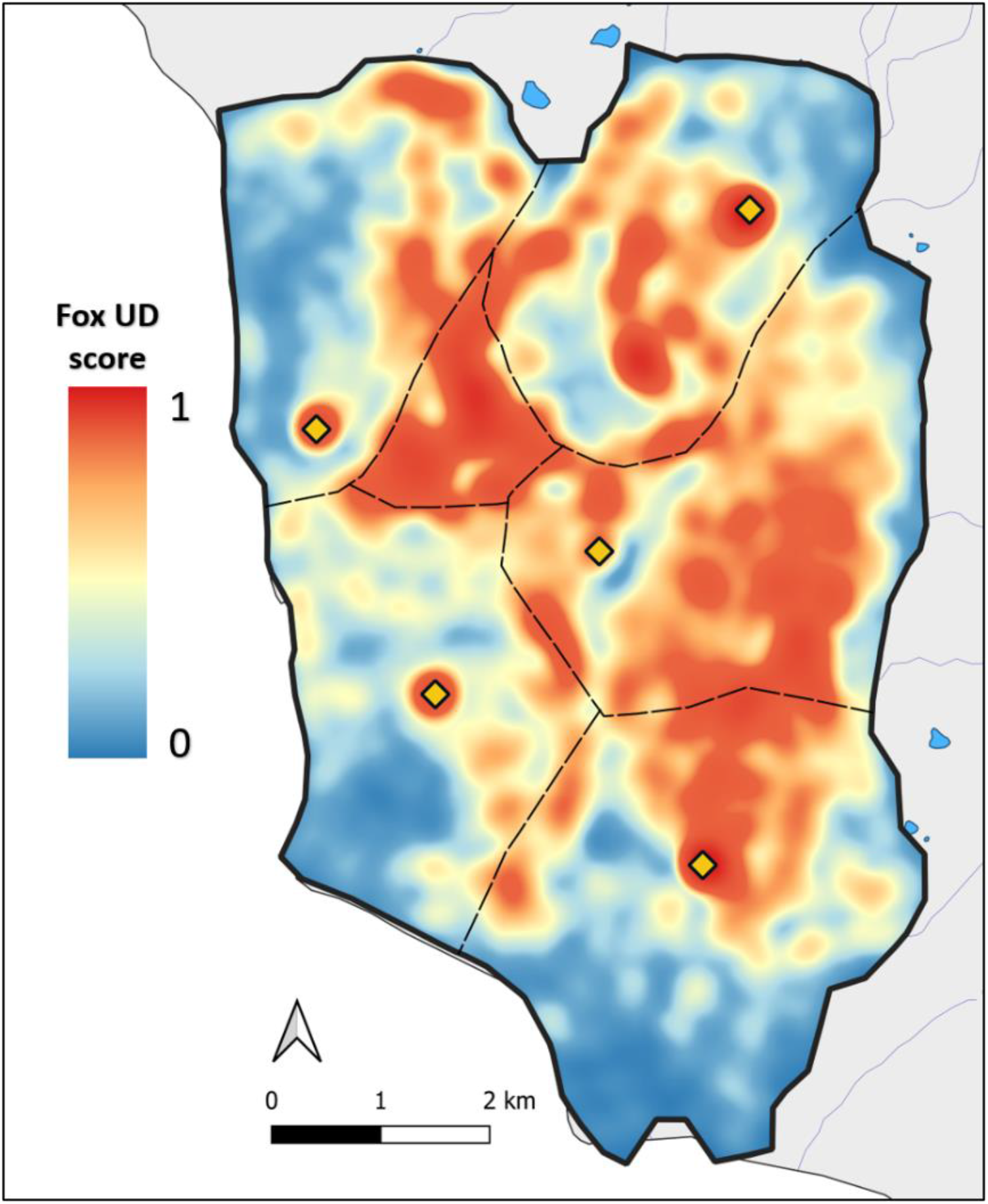
Arctic fox activity landscape generated from 20,961 GPS locations classified in the active state by a hidden Markov model. The activity landscape reflects fox utilization distribution (UD) based on data from 13 individuals living in 6 territories, tracked during 10 days at the end of June 2019 on Bylot Island. A UD smoothing parameter of 300 m was used to generate this activity landscape (see Appendix S1: Fig. S4 for activity landscapes generated from other smoothing parameters). The color scale reflects fox UD score (from 0 to 1) and thus probability of presence of a fox. Yellow diamonds locate the 5 reproductive dens, dotted lines identify the approximate boundaries of fox pair territories, and the thick black line is the contour of the study area.

### Probability of predation of baits

The artificial prey experiment showed high adequacy between the predator activity landscape and the predation risk landscape. Baits were more likely to be consumed where fox UD score was high (Table 1, Fig. 3a), whatever the UD smoothing parameter (Table 1).

**Table 1.** Results from binomial mixed models testing the effect of fox UD score on the probability of predation of baits, with patch ID nested in territory ID fitted as random effects, for the 5 UDs with smoothing parameter ranging from 200 to 400 m (n = 428 baits). See Appendix S1: Table S2 for variance values of random effects.

| UD smoothing parameter (m) | Fixed effect | Estimate [95% CI] | z value | p value |
| --- | --- | --- | --- | --- |
| 200 | (Intercept) | -0.25 [-1.17, 0.68] | -0.60 | 0.546 |
|  | Fox UD score | 3.56 [-0.02, 7.45] | 1.91 | 0.056 |
| 250 | (Intercept) | -0.35 [-1.29, 0.60] | -0.82 | 0.414 |
|  | Fox UD score | 3.28 [0.31, 6.49] | 2.12 | 0.034 |
| 300 | (Intercept) | -0.48 [-1.44, 0.49] | -1.07 | 0.284 |
|  | Fox UD score | 3.24 [0.59, 6.09] | 2.36 | 0.013 |
| 350 | (Intercept) | -0.61 [-1.61, 0.38] | -1.31 | 0.191 |
|  | Fox UD score | 3.23 [0.79, 5.84] | 2.56 | 0.011 |
| 400 | (Intercept) | -0.73 [-1.77, 0.29] | -1.50 | 0.133 |
|  | Fox UD score | 3.18 [0.89, 5.62] | 2.69 | 0.007 |

**Fig. 3.**
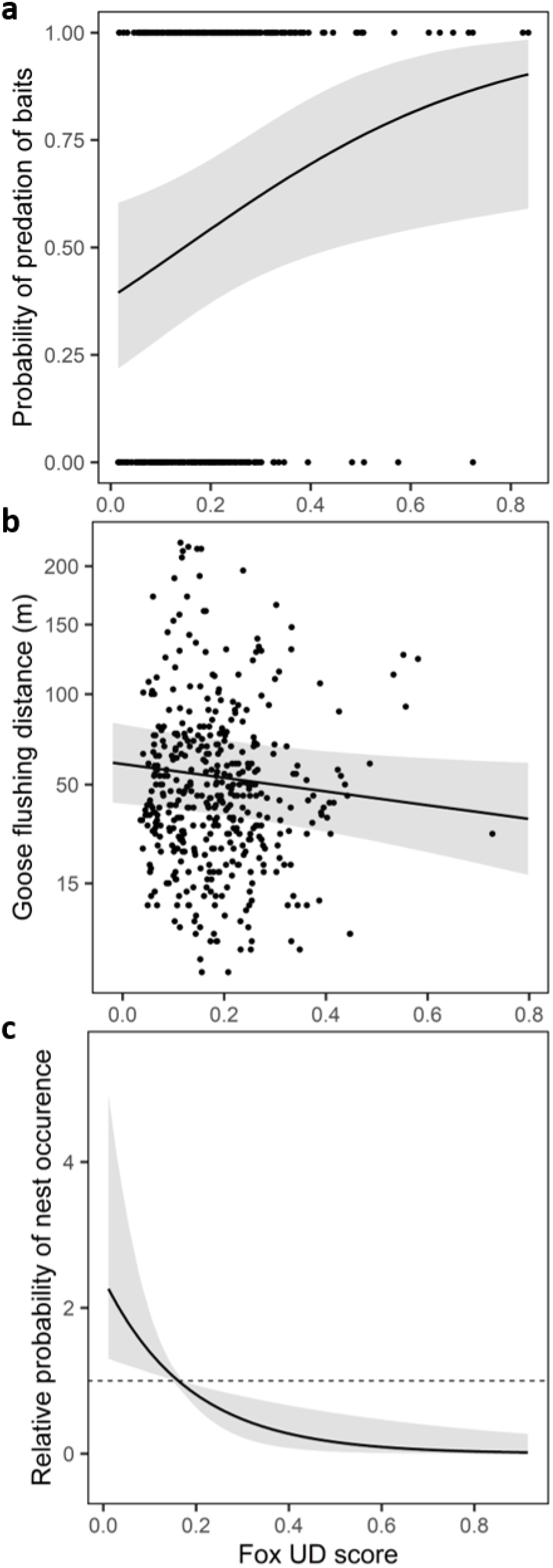
Predicted effect of fox UD score on (a) probability of predation of baits (0 = bait not eaten, 1 = bait eaten, n = 428), (b) goose flushing distance (n = 458) and (c) relative probability of occurrence of nests from birds nesting in habitats easily accessible to foxes (n = 44 nests from 10 species). In (b), we back-transformed goose flushing distance and fox UD score before plotting (goose flushing distance had been square-root transformed and fox UD score had been centered and standardized in linear models), leading to irregular y-axis increments. In (c) the dashed horizontal line represents a relative probability of occurrence of 1, with values below and above 1 indicating lower and higher probabilities of occurrence than random, respectively. The gray area represents the 95% confidence interval of (a) the fitted logistic regression, (b) the linear regression and (c) the relative probability of occurrence obtained by bootstrap. For these representations we used fox UD scores generated with an intermediate smoothing parameter of 300 m, and (c) nest areas generated with an intermediate radius of 2000 m.

### Snow goose nest defense behavior

Snow geese showed higher level of nest defense when nesting in areas of high predation risk, as shown by the negative relationship between flushing distance and fox UD score (Table 2, Fig. 3b). Although slope estimates were consistently negative for the 5 UDs, the slope lessened and lost its significance as the UD smoothing parameter increased. Geese also showed a weaker level of nest defense when they had a relatively small clutch and saw the observer approaching from far away, as shown by the significant effects of clutch size and starting distance on flushing distance (no effect of smoothing parameter, Table 2). Flushing distance did not vary with observation date.

**Table 2.**
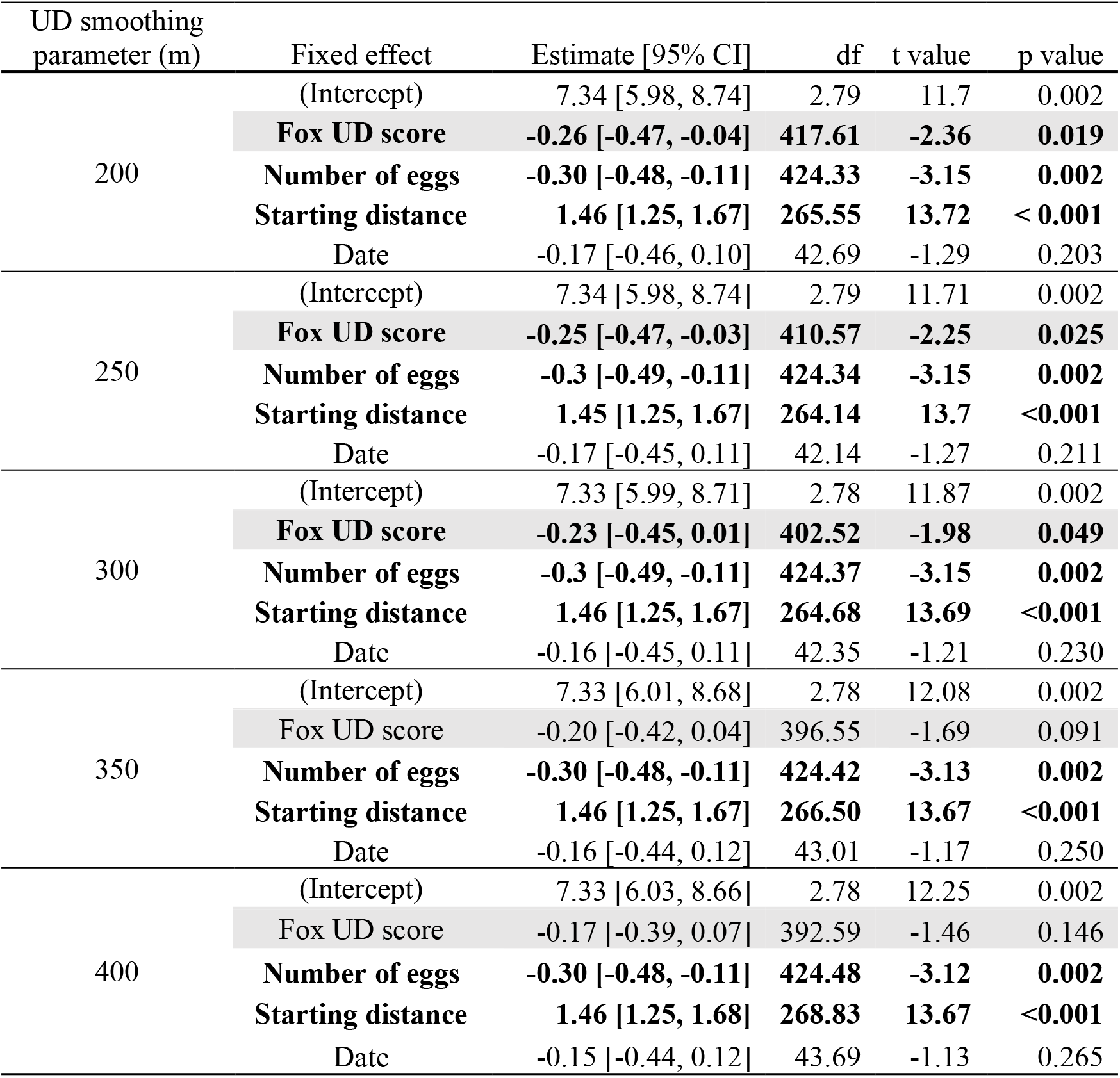
Results from linear mixed models testing the effect of fox UD score on goose flushing distance. Number of eggs, starting distance of the observer and date were included as covariates. Territory ID and observer ID were fitted as random effects and models were repeated for the 5 UDs with smoothing parameter ranging from 200 to 400 m (n = 458 goose nests). All fixed effects are standardized and the effect of interest (fox UD score) is highlighted in gray. Significant effects are in bold. See Appendix S1: Table S3 for variance values of random effects.

### Nest distribution of incidental prey

For bird species nesting in habitats easily accessible to foxes, nests were more likely to occur where fox UD score was low, compared to random locations (Table 3, Fig. 3c). Specifically, the probability of nest occurrence was approximately twice higher than the probability of occurrence of a random location where fox UD score was the lowest, and the probability of nest occurrence also declined steeply as fox presence increased (Fig. 3c). The effect of fox UD score on nest distribution was significant or almost significant (with p values only slightly over 0.05 and the upper limit of confidence intervals slightly over 0) for all 5 fox activity landscapes (smoothing parameters of 200–400 m) and 5 nest area sizes (radii of 1000–3000 m) (Table 3).

**Table 3.** Results from conditional logistic regressions with a use-available design testing the effect of fox UD score on the nest distribution of (1) birds nesting in habitats easily accessible to foxes (n = 44 nests from 10 species) and (2) birds nesting in micro-habitats providing a refuge against foxes (n = 65 nests from 3 species). The fox UD score of each nest located in an easily accessible habitat was compared to the fox UD scores of 50 random locations surrounding the nest within a predetermined nest area. The fox UD score of each nest located in a refuge was compared to the fox UD score of ≤ 50 islets surrounding the nest within the nest area. Coefficient estimates are presented for 25 models, each reflecting a given size of the nest area (from 1000 to 3000 m) and UD smoothing parameter (from 200 to 400 m). Significant effects are in bold.

| Radius of nest area (m) | UD smoothing parameter (m) | (1) Nests in easily accessible habitats |  |  | (2) Nests in refuges |  |  |
| --- | --- | --- | --- | --- | --- | --- | --- |
|  |  | Coefficient [95% CI] | z value | p value | Coefficient [95% CI] | z value | p value |
| 1000 | 200 | **-9.81 [-18.42, -1.20]** | **-2.23** | **0.026** | 0.93 [-7.36, 9.23] | 0.22 | 0.826 |
|  | 250 | **-7.12 [-13.86, -0.39]** | **-2.07** | **0.038** | 0.50 [-6.14, 7.13] | 0.15 | 0.884 |
|  | 300 | **-5.58 [-11.17, -0.002]** | **-1.96** | **0.050** | 1.13 [-5.45, 5.70] | 0.04 | 0.965 |
|  | 350 | -4.46 [-9.23, 0.32] | -1.83 | 0.067 | -0.20 [-5.06, 4.67] | -0.08 | 0.937 |
|  | 400 | -3.53 [-7.69, 0.63] | -1.66 | 0.097 | -0.46 [-4.84, 3.92] | -0.20 | 0.838 |
| 1500 | 200 | **-10.61 [-18.60, -2.61]** | **-2.60** | **0.009** | 4.43 [-1.66, 10.52] | 1.43 | 0.154 |
|  | 250 | **-7.70 [-13.82, -1.57]** | **-2.46** | **0.014** | 3.24 [-1.47, 7.96] | 1.35 | 0.178 |
|  | 300 | **-6.04 [-11.05, -1.04]** | **-2.37** | **0.018** | 2.40 [-1.49, 6.30] | 1.21 | 0.226 |
|  | 350 | **-4.89 [-9.13, -0.66]** | **-2.26** | **0.024** | 1.79 [-1.54, 5.12] | 1.05 | 0.293 |
|  | 400 | **-4.01 [-7.68, -0.34]** | **-2.14** | **0.032** | 1.36 [-1.58, 4.31] | 0.91 | 0.365 |
| 2000 | 200 | **-9.87 [-17.33, -2.41]** | **-2.59** | **0.009** | 3.50 [-1.79, 8.78] | 1.30 | 0.195 |
|  | 250 | **-7.04 [-12.67, -1.41]** | **-2.45** | **0.014** | 2.44 [-1.64, 6.51] | 1.17 | 0.241 |
|  | 300 | **-5.42 [-9.95, -0.88]** | **-2.34** | **0.019** | 1.74 [-1.63, 5.11] | 1.01 | 0.312 |
|  | 350 | **-4.29 [-8.07, -0.50]** | **-2.22** | **0.026** | 1.28 [-1.60, 4.16] | 0.87 | 0.385 |
|  | 400 | **-3.44 [-6.68, -0.21]** | **-2.09** | **0.037** | 0.97 [-1.56, 3.50] | 0.75 | 0.453 |
| 2500 | 200 | **-8.05 [-14.82, -1.28]** | **-2.33** | **0.020** | 3.47 [-1.82, 8.76] | 1.29 | 0.199 |
|  | 250 | **-5.68 [-10.77, -0.59]** | **-2.19** | **0.029** | 2.55 [-1.51, 6.60] | 1.23 | 0.218 |
|  | 300 | **-4.35 [-8.44, -0.26]** | **-2.08** | **0.037** | 1.92 [-1.41, 5.25] | 1.13 | 0.258 |
|  | 350 | **-3.45 [-6.86, -0.04]** | **-1.98** | **0.048** | 1.47 [-1.38, 4.31] | 1.01 | 0.312 |
|  | 400 | -2.78 [-5.69, 0.14] | -1.87 | 0.062 | 1.15 [-1.36, 3.64] | 0.90 | 0.371 |
| 3000 | 200 | **-7.03 [-13.61, -0.45]** | **-2.10** | **0.036** | 3.78 [-1.48, 9.05] | 1.41 | 0.159 |
|  | 250 | **-4.99 [-9.90, -0.08]** | **-1.99** | **0.047** | 2.79 [-1.21, 6.80] | 1.37 | 0.171 |
|  | 300 | -3.83 [-7.75, 0.09] | -1.91 | 0.056 | 2.16 [-1.11, 5.43] | 1.30 | 0.195 |
|  | 350 | -3.02 [-6.28, 0.23] | -1.82 | 0.068 | 1.70 [-1.07, 4.48] | 1.20 | 0.229 |
|  | 400 | -2.42 [-5.19, 0.35] | -1.71 | 0.087 | 1.37 [-1.06, 3.80] | 1.11 | 0.269 |

For bird species nesting in refuges, fox UD score did not affect the probability of nest occurrence, whatever the smoothing parameter or nest area radius (Table 3). Fox UD scores of nesting locations were not statistically different from those of random islets (Table 3). Variation in the number of random islets available for testing did not affect results (Appendix S2).

## Discussion

Considering simultaneously all actors interacting in a heterogeneous landscape is needed to fully assess the ecological context of the landscape of fear and its consequences on natural communities (Gaynor et al. 2019). Using high resolution arctic fox GPS data, behavioral observations and field experiments, we demonstrated that fine scale variation in space use of active predators accurately reflects spatial variation in predation risk, and explains anti-predator behavior of a main prey and nest distribution of some incidental prey species in an Arctic terrestrial community (Fig. 4). Overall, our study demonstrates the impacts of predator activity on the behavioral decisions and distribution of prey and highlights the spatial dimension of predator-prey relationships.

**Fig. 4.**
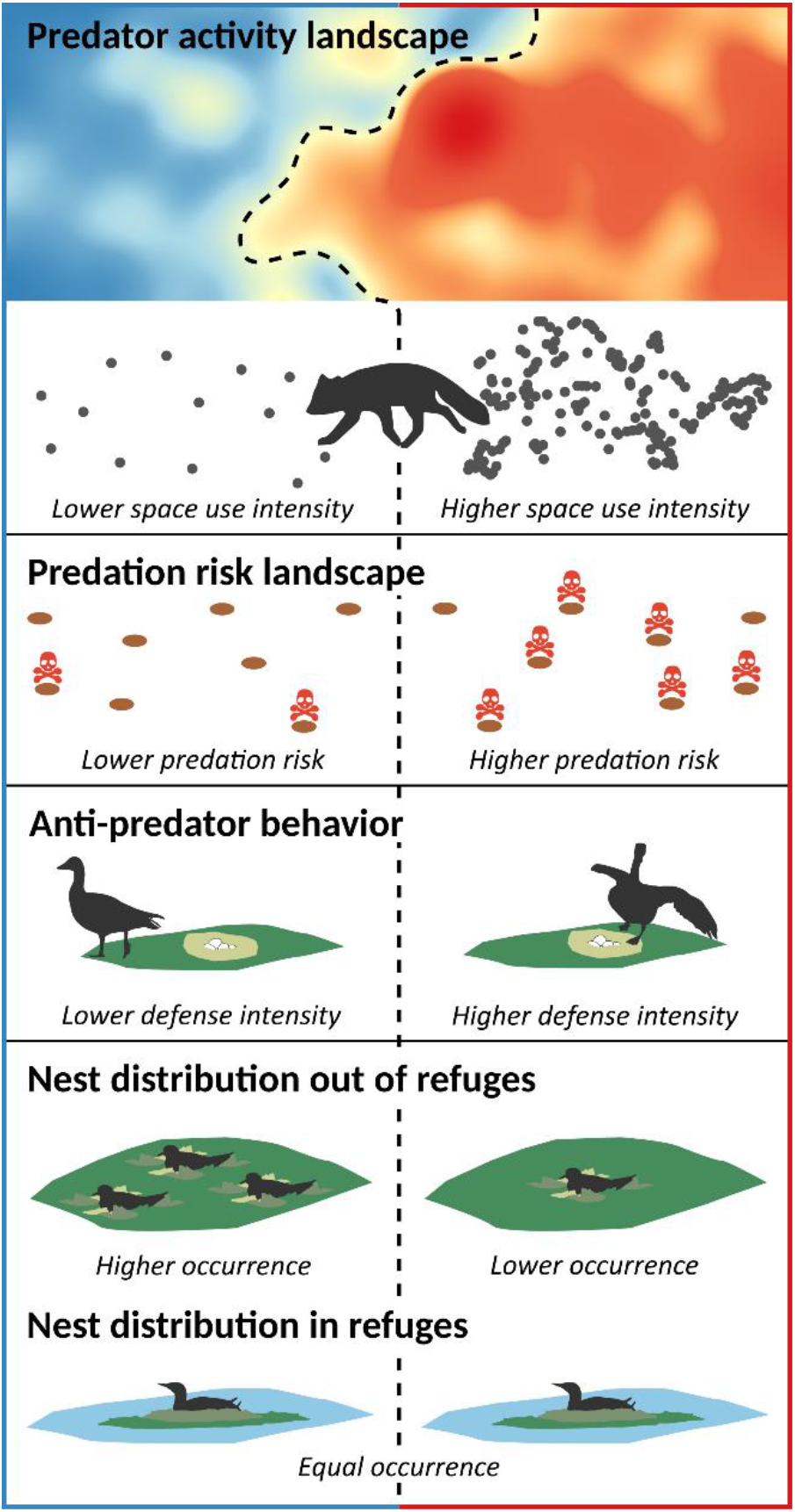
Landscape of fear context in a terrestrial Arctic community. The predator activity landscape generates a landscape of predation risk and predicts anti-predator response and distribution of prey. The illustrated predator activity landscape shows the multiple spatial gradients of intensity in arctic fox space use (low in blue, high in red). Causation between the predator activity landscape and the predation risk landscape is demonstrated by an artificial prey experiment. The predator activity landscape predicts anti-predator behavior of a main prey (here, snow goose) and nest distribution of incidental prey (here, a shorebird) nesting in habitats accessible to foxes. However, the nest distribution of incidental prey nesting in predation refuges such as islets (here, a loon) is independent of the predator activity landscape at its measured spatial resolution.

### The predator activity landscape as a predation risk landscape

The use of fox movement data collected at a high frequency, in combination with the identification of active and resting behavioral states yielded a predator activity landscape that robustly depicted fine scale variation in fox intensity of space use in our study area. An artificial prey experiment using baits demonstrated that the probability of predation was higher in areas highly used by foxes, and therefore that spatial variation in fox space use was related to predation risk for prey. Our sensitivity analyses also confirmed the robustness of our results, which were consistent across our range of UD smoothing parameters (Tables 1, 2, 3). Most importantly, when the adequacy between the predator activity landscape and the predation risk landscape is demonstrated, the predator activity landscape allows to identify predator “hotspots” where risk of predation should be the highest, and thus creates opportunities to better understand the spatial dimension of predator-prey interactions.

Furthermore, to obtain a good estimation of the probability of predator-prey encounter and thus predation risk, it is crucial to accurately model spatial variation in predator space use intensity (or predator density), which is not always possible using proxies of predator space use (e.g., habitat features), or when only a limited number of predator locations are used. Also, prey in multi-predator systems may have to deal with multiple and contrasting landscapes of risk (Thaker et al. 2011, Gaynor et al. 2019). In our system, even though foxes are the main predators of all nesting prey (Gauthier et al. 2011), the occasional presence of territorial ermines (*Mustela erminea*) in years of high lemming abundance may increase predation risk in some areas, while nesting association with snowy owls may reduce predation risk (Bêty et al. 2001). Our predator activity landscape accurately depicted spatial variation in predation risk as we were able to collar all foxes living in our study area, where neither ermines nor snowy owls were detected.

### Goose nest defense intensity partly explained by the predator activity landscape

It is difficult for primary prey species to reduce their exposure to predation risk because they are actively searched by predators. As such, highly conspicuous nesting snow geese can hardly use spatial avoidance to reduce predation risk as active foxes select patches where goose nest density is highest (Grenier-Potvin et al. 2020). Geese can nonetheless adopt a fine scale response by using nest defense when predation risk of their nest is imminent (Bêty et al. 2002, Lima 2009). We indeed found that snow geese nesting in areas highly used by foxes showed a higher level of nest defense compared to geese nesting in less used areas. This likely results from plastic adjustments of anti-predator behavior in response to variation in predation risk, such as female ungulates showing greater levels of vigilance when foraging in habitats associated to higher predation risk caused by wolf presence (Laundré et al. 2001). Assessing anti-predator behavior on the same individuals along a gradient of predation risk (Fontaine and Martin 2006, Mathot et al. 2011) would be required to fully understand the underlying mechanisms explaining the relationship between snow geese nest defence and the predator activity landscape. Nonetheless, our results suggest that predator space use impacts large prey behavior, which ultimately imposes costs on their fitness (Cresswell 2008).

The effect of the fox activity landscape on goose flushing distances was moderate, and model outputs slightly differed according to UD smoothing parameters (Table 2). Variables affecting goose nest defense other than those considered in this study, like the parent’s physiological state or the presence of conspecifics (Kazama et al. 2011), may further explain variation in goose nest defense. Also, although we limited effects of the incubation stage of nest defense intensity by approaching nests during a short period of time and by controlling potential effects of date in our models, nesting asynchrony may have created uncontrolled variation in nest defense intensity.

Finally, aspects of the physical landscape that affect nest visibility and thus predator detection may affect how prey perceive the level of predation risk and respond to the predator activity landscape (Gaynor et al. 2019).

### Habitat structure modulates the effect of predator activity landscape on incidental prey

As proposed by Gaynor et al (2019), we found evidence that the physical landscape can intervene in the ecological context of the landscape of fear. Indeed, cackling geese, glaucous gulls and red-throated loons nest on islets that are poorly accessible to foxes and serve as refuges (Gauthier et al. 2015). Hence, species using such structures can better afford to have their nest surrounded by a relatively risky landscape, explaining why they did not respond to the fox activity landscape at its measured spatial resolution. This is not the case for species nesting in adjacent habitats easily accessible to foxes. Indeed, the probability of occurrence of these other nesting birds was lower in areas heavily used by foxes. These species may perceive predation risk and avoid nesting in areas highly used by foxes, as birds can shift nest location when encountering predators during nest building (Peluc et al. 2008). Selection of safe habitats may also be fixed genetically through selection. For example, some shorebirds nest strictly on stony riverbanks (Léandri-Breton and Bêty 2020), a habitat avoided by foxes (Grenier-Potvin et al. 2020). How habitat structure modulates the predator activity landscape and its effects on incidental prey is a rich topic for research on predator-mediated interactions between prey species.

The relationship between the predator activity landscape and nest distribution may also result from consumptive effects of predation, as nests located in areas highly used by foxes may have been preyed upon before being detected by observers. Monitoring fine-scale bird movements during nest establishment (Gilbert et al. 2016) and determining nest location prior to any nest predation event should help differentiating the roles of consumptive and non-consumptive effects of predation, and to investigate the ability of nesting birds to perceive and respond to predation risk.

Finally, animals face a variety of physiological, phylogenetic or ecological constraints that limit their ability to assess predation risk and respond to the landscape of fear (Jordan and Ryan 2015, Gaynor et al. 2019). Measuring the landscape of fear directly, by assessing how prey perceive predation risk, would increase our understanding of complex relationships between predation risk and prey responses, despite the challenges that this approach entails (Gaynor et al. 2019).

## Conclusion

Our study demonstrates the value of using fine scale predator movements to characterize the landscape of predation risk in landscape of fear studies. It also highlights how the predator activity landscape influences the anti-predator behavior of a main prey and the nest distribution of incidental prey species from different guilds. Assessing the effects of the landscape of fear in a community allows to better understand prey species distribution and behavior, providing insights on the structure and functioning of a community.

## Acknowledgments

We thank M.-P. Poulin, R. Gravel, C. Chevallier, S. Lai, G. Roy, M. Loyer, F. Letourneux, M.Z. Corbeil-Robitaille and A. Florea for their ideas and field work. Research support was received from Natural Sciences and Engineering Research Council of Canada (NSERC), Canada Foundation for Innovation, Canada Research Chairs Program, Network of Centers of Excellence of Canada ArcticNet, Northern Scientific Training Program (Polar Knowledge Canada), Parks Canada Agency and Polar Continental Shelf Program (Natural Resources Canada). J. Clermont received a Weston Family Awards in Northern Research and scholarships from NSERC and Fonds de Recherche du Québec.

J. Clermont, J. Bêty and D. Berteaux conceived the study with contribution from coauthors. All authors contributed to field planning or data acquisition. J. Clermont conducted statistical analyses, with significant help from A. Grenier-Potvin and contribution from coauthors. J. Clermont led the writing, with contribution from coauthors.

## Appendix S1 Supplementary tables and figures

**Table S1.**
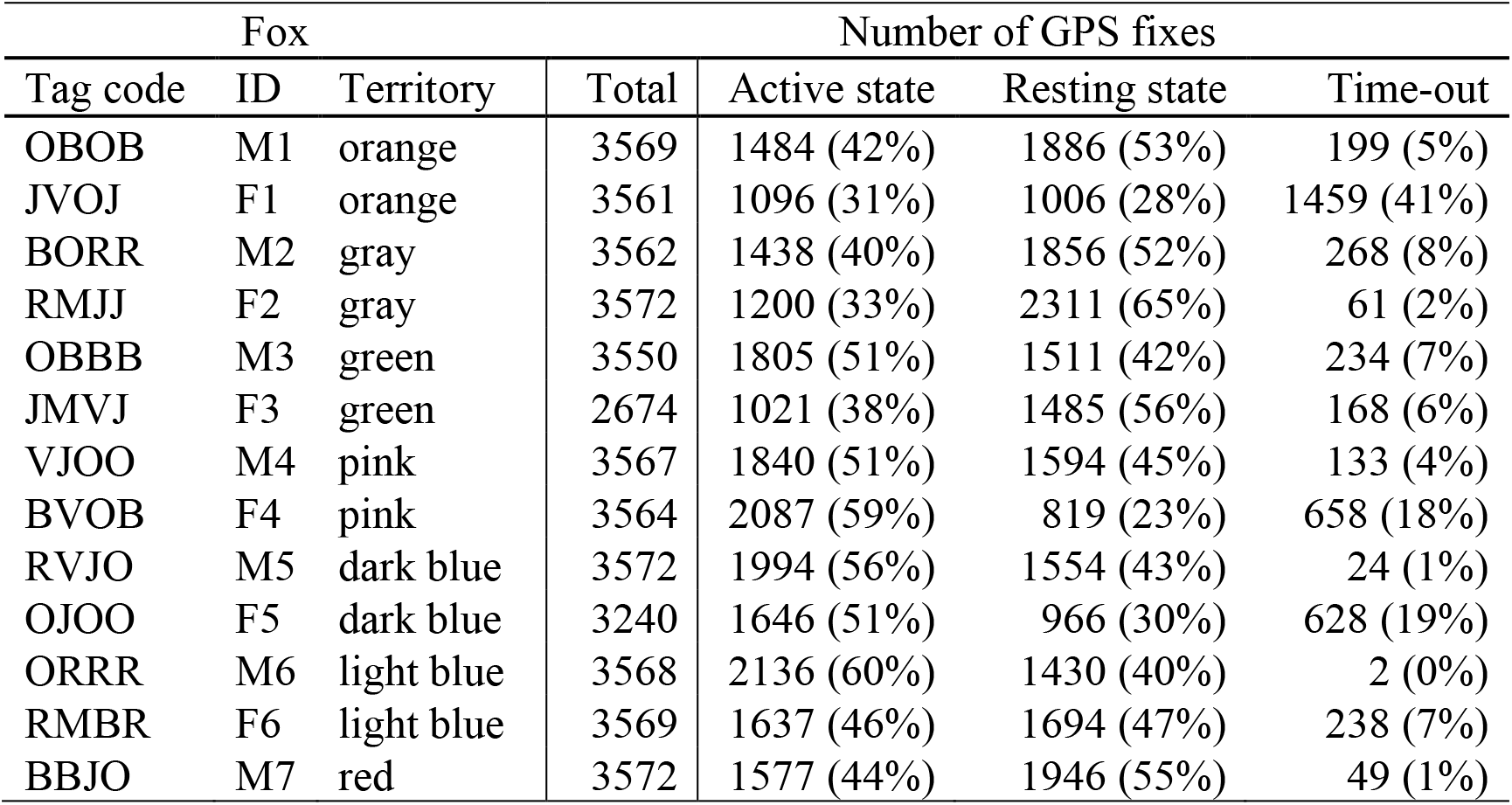
Number of GPS fixes obtained for 13 arctic foxes tracked during 10 days at the end of June 2019 on Bylot Island (Nunavut, Canada). The tag code, ID (also indicating sex, M: male, F: female) and territory color (see Fig. 1) are given for each fox. GPS fixes were assigned an active or resting state using a hidden Markov model (HMM). Fixes with unknown location (time-outs) due to missing connection with satellites, which mostly occurred when foxes were inside their den, were excluded from analyses. The number of fixes per individual was nearly 3600 (10 days at a 4-min fix interval) for all foxes except F3 (due to a battery failure).

**Table S2.** Variance values for random effects of binomial mixed models testing the effect of fox UD score on the probability of predation of baits, with patch ID nested in territory ID fitted as random effects, for the 5 UDs with smoothing parameter ranging from 200 to 400 m (n = 428 baits).

| UD smoothing parameter (m) | Random effect | Variance | SD |
| --- | --- | --- | --- |
| 200 | Patch: territory | 0.40 | 0.63 |
|  | Territory | 0.65 | 0.81 |
| 250 | Patch: territory | 0.39 | 0.63 |
|  | Territory | 0.64 | 0.80 |
| 300 | Patch: territory | 0.38 | 0.62 |
|  | Territory | 0.64 | 0.80 |
| 350 | Patch: territory | 0.37 | 0.61 |
|  | Territory | 0.64 | 0.80 |
| 400 | Patch: territory | 0.37 | 0.61 |
|  | Territory | 0.65 | 0.81 |

**Table S3.** Variance values for random effects of linear mixed models testing the effect of fox UD score on goose flushing distance, with territory ID and observer ID fitted as random effects, for the 5 UDs with smoothing parameter ranging from 200 to 400 m (n = 458 goose nests).

| UD smoothing parameter (m) | Random effect | Variance | SD |
| --- | --- | --- | --- |
| 200 | Territory | 1.44 | 1.20 |
|  | Observer | 0.10 | 0.32 |
|  | Residuals | 3.78 | 1.94 |
| 250 | Territory | 1.44 | 1.20 |
|  | Observer | 0.10 | 0.31 |
|  | Residuals | 1.38 | 1.94 |
| 300 | Territory | 1.39 | 1.18 |
|  | Observer | 0.10 | 0.31 |
|  | Residuals | 3.79 | 1.95 |
| 350 | Territory | 1.34 | 1.16 |
|  | Observer | 0.10 | 0.32 |
|  | Residuals | 3.80 | 1.95 |
| 400 | Territory | 1.29 | 1.14 |
|  | Observer | 0.10 | 0.32 |
|  | Residuals | 3.81 | 1.95 |

**Fig. S1.**
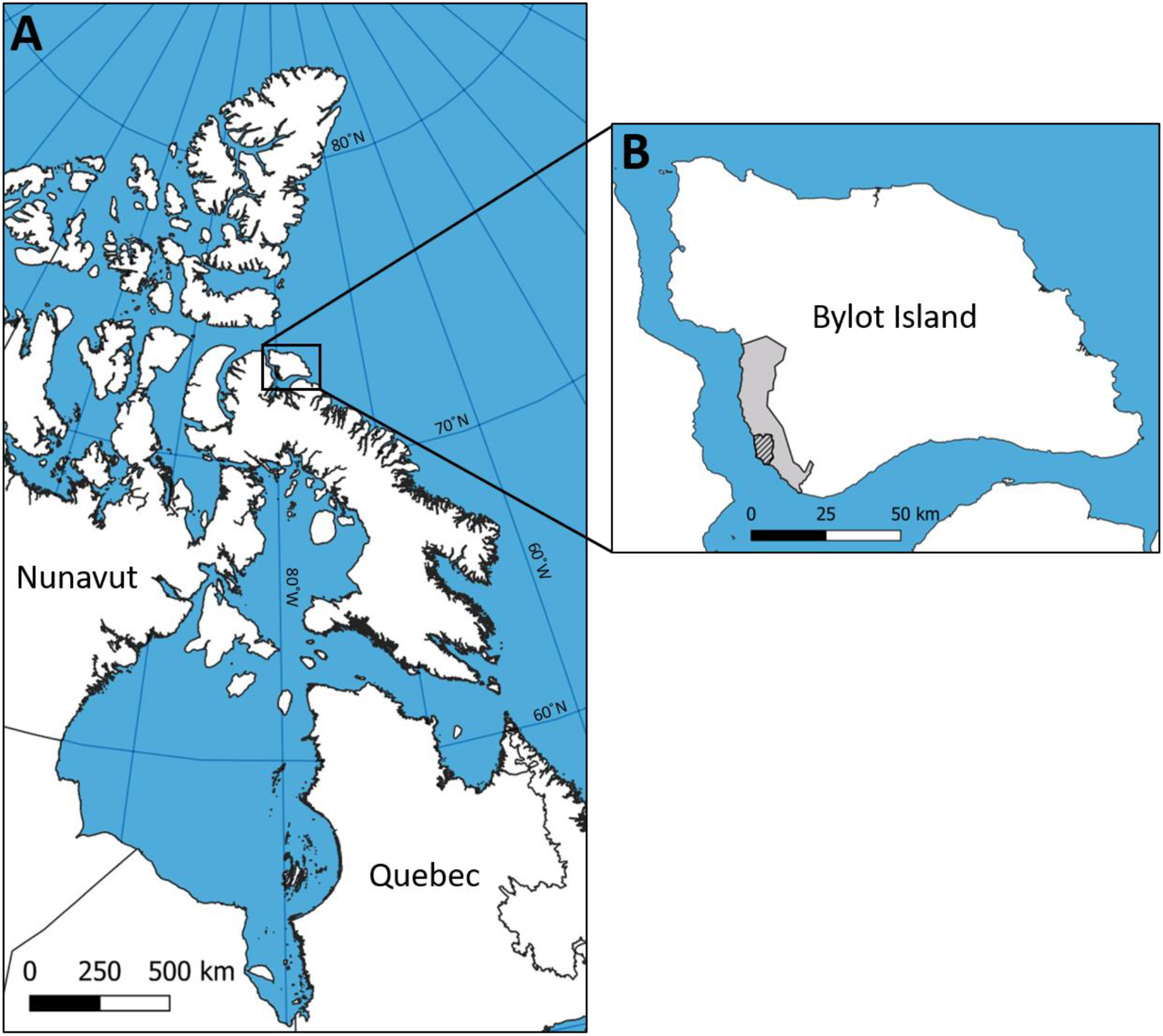
Geographical context of the study area. Panel A locates Bylot Island (72°53’ N, 79°54’ W) in Nunavut, Canada, which is enlarged in B. The hatched area depicts our study area containing 6 arctic fox territories used by 6 fox pairs and one additional individual (Fig. 1), while the larger gray area depicts Bylot Island’s entire field site.

**Fig. S2.**
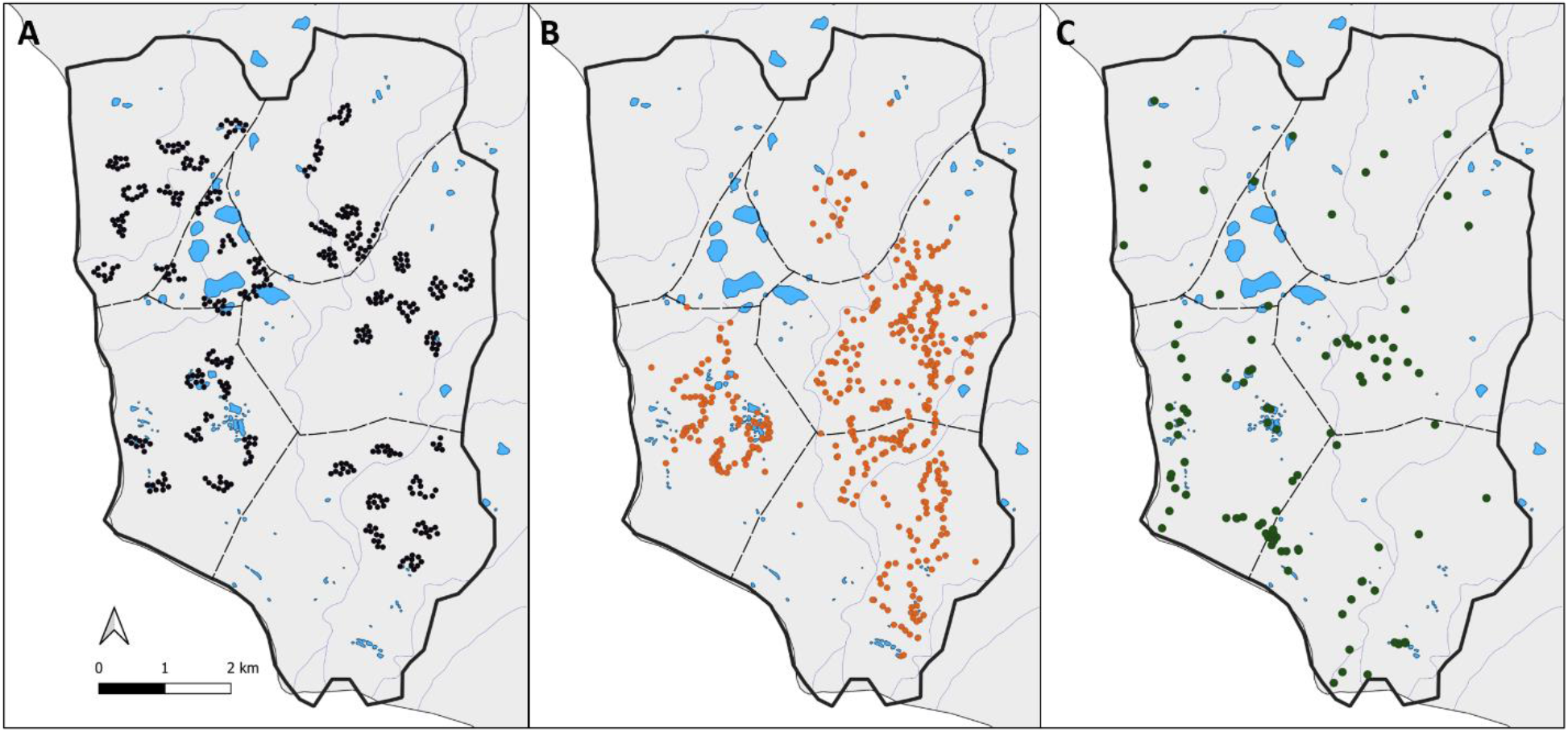
Distribution of A) fox baits used in the artificial prey experiment (n = 428, black dots), nests used to evaluate snow goose anti-predator behavior (n = 458, orange dots), and C) nests of fox incidental prey, that is birds other than snow geese (n = 109, green dots). The dotted lines show the approximate boundaries of fox pair territories while the thick black line is the contour of the study area. Lakes and large ponds are in blue.

**Fig. S3.**
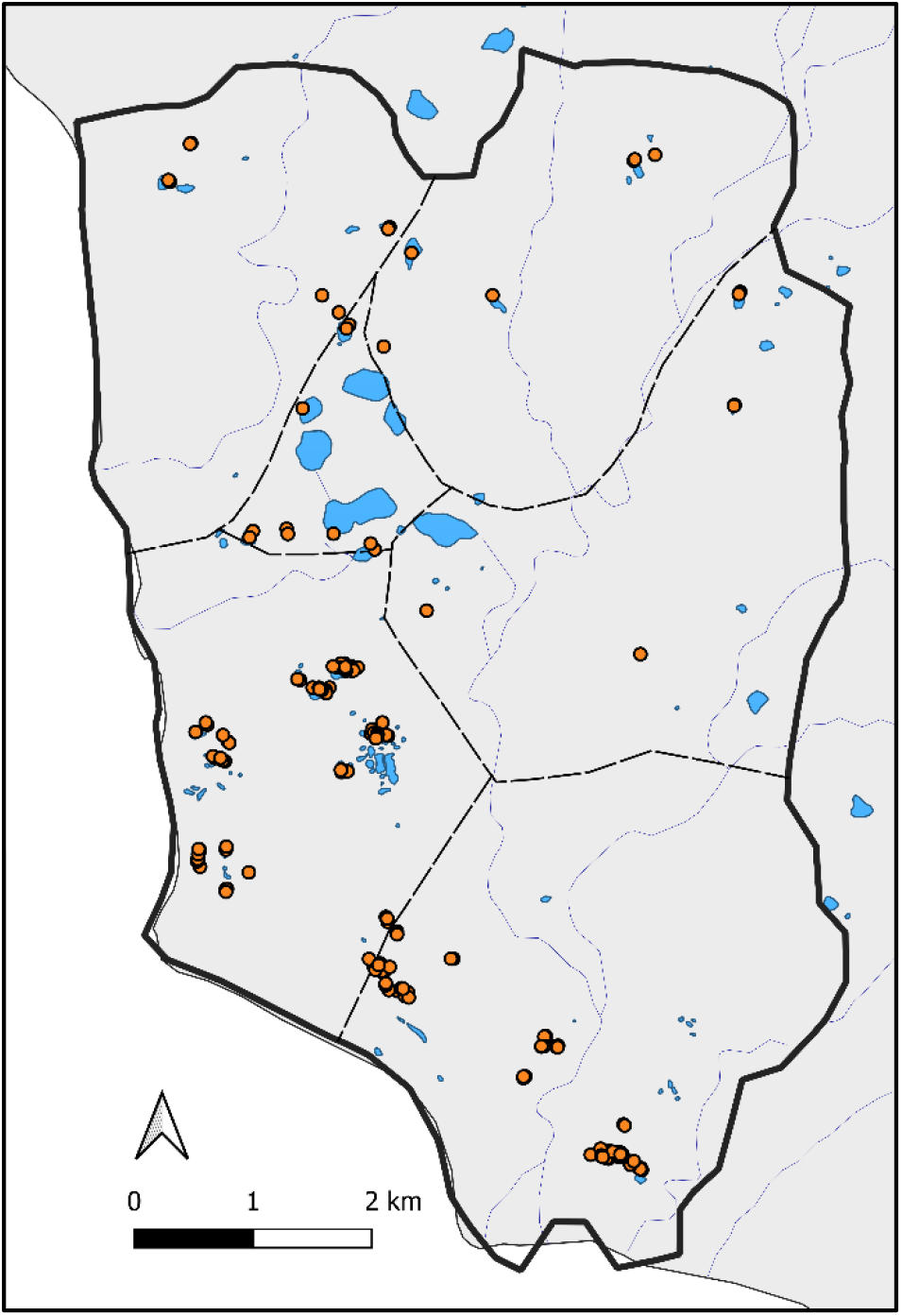
Distribution of the 377 islets (orange dots) located in ponds, lakes and wetlands. Many dots are superimposed at this spatial scale. The dotted lines show the approximate boundaries of fox pair territories while the thick black line is the contour of the study area. Lakes and large ponds are in blue.

**Fig. S4.**
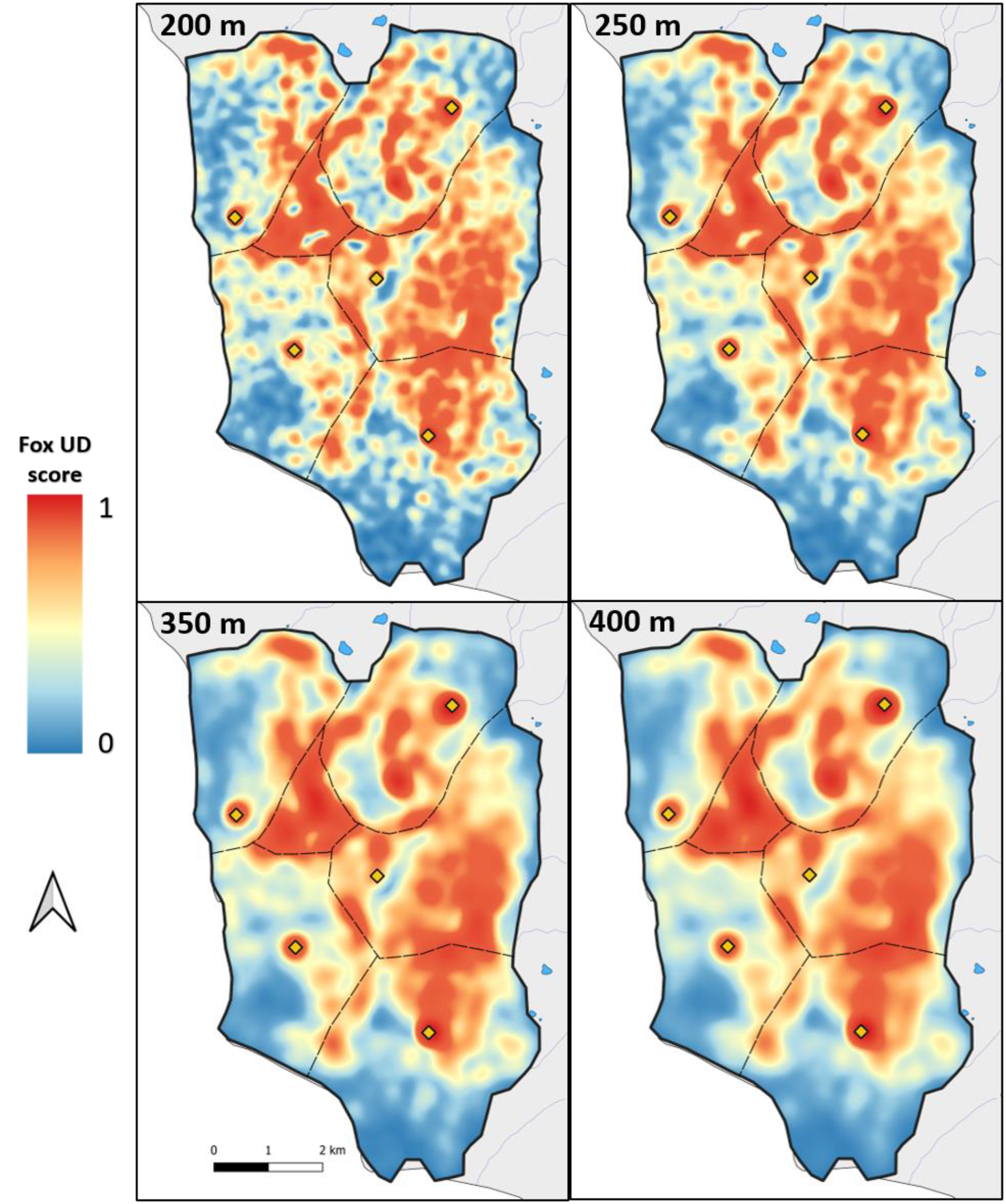
Arctic fox activity landscapes generated from 20,961 GPS locations classified in the active state by a hidden Markov model, using UD smoothing parameters ranging from 200 to 400 m, as indicated on the top left corner of each map (see Fig. 2 in the main text for activity landscape using the intermediate smoothing parameter of 300 m). The activity landscape reflects fox utilization distribution (UD) based on data from 13 individuals living in 6 territories during 10 days at the end of June 2019 on Bylot Island. The color scale reflects fox UD score (from 0 to and thus probability of presence of a fox. Yellow diamonds locate the 5 reproductive dens, dotted lines identify the approximate boundaries of fox pair territories, and the thick black line is the contour of the study area.

## Appendix S2 Complementary analyses for distribution of birds nesting in refuges

The number of available islets per nest varied according to the radius of the nest area, with 26 ± 12 (range 3–50) available islets for a radius of 1000 m, 40 ± 12 (3–50) for 1500 m, 46 ± 10 (3–50) for 2000 m, 47 ± 9 (9–50) for 1500 m and 48 ± 7 (10–50) for 3000 m. Results from these models are presented in Table 3 (main text).

We conducted two complementary analyses to verify whether using an unbalanced number of available locations affected results. First, we compared fox UD scores at bird nests to fox UD scores at available islets as described in Methods (3-Nest distribution of incidental prey/Second set of models), but using more balanced sample sizes of ≤ 10 islets instead of ≤ 50. We could not use a totally balanced design as the minimum number of islets was only 3. Obtained results (Table S1) did not differ from those presented in the main text (Table 3).

Second, we compared fox UD scores at bird nests to fox UD scores at available locations as described in Methods (3-Nest distribution of incidental prey/First set of models), thus comparing UD scores of used islets to 50 random locations picked anywhere in the nest area, excluding water bodies (this yielded a balanced design). An advantage of this approach is that it also allowed us to verify whether drawing available locations from georeferenced islets rather than anywhere in the nest area affected our conclusions. Results (Table S2) did not differ from those presented in the main text (Table 3).

**Table S1.** Results from conditional logistic regressions with a use-available design testing the effect of fox UD score on the nest distribution of birds that nest in micro-habitats providing partial refuge against foxes (n = 65 nests from 3 species). The fox UD score of each nest was compared to the fox UD scores of 50 random locations surrounding the nest within the nest area. Coefficient estimates are presented for 25 models, each reflecting a given size of the nest area (from 1000 to 3000 m) and UD smoothing parameter (from 200 to 400 m).

| Radius of nest area (m) | UD smoothing parameter (m) | Coefficient [95% CI] | z value | p value |
| --- | --- | --- | --- | --- |
| 1000 | 200 | 0.41 [-7.95, 8.76] | 0.10 | 0.924 |
|  | 250 | 0.32 [-6.37, 7.01] | 0.09 | 0.926 |
|  | 300 | 0.18 [-5.47, 5.82] | 0.06 | 0.952 |
|  | 350 | -0.02 [-4.97, 4.94] | -0.01 | 0.995 |
|  | 400 | -0.21 [-4.70, 4.27] | -0.09 | 0.926 |
| 1500 | 200 | 2.04 [-4.16, 8.24] | 0.64 | 0.519 |
|  | 250 | 1.41 [-3.40, 6.21] | 0.57 | 0.566 |
|  | 300 | 0.89 [-3.06, 4.85] | 0.44 | 0.658 |
|  | 350 | 0.50 [-2.87, 3.89] | 0.29 | 0.771 |
|  | 400 | 0.24 [-2.77, 3.24] | 0.15 | 0.878 |
| 2000 | 200 | 4.12 [-1.67, 9.91] | 1.40 | 0.163 |
|  | 250 | 2.90 [-1.54, 7.33] | 1.28 | 0.201 |
|  | 300 | 2.06 [-1.59, 5.70] | 1.11 | 0.269 |
|  | 350 | 1.52 [-1.59, 4.63] | 0.96 | 0.338 |
|  | 400 | 1.20 [-1.53, 3.93] | 0.86 | 0.390 |
| 2500 | 200 | 3.75 [-1.88, 9.39] | 1.31 | 0.192 |
|  | 250 | 2.74 [-1.56, 7.05] | 1.25 | 0.211 |
|  | 300 | 2.10 [-1.41, 5.62] | 1.17 | 0.241 |
|  | 350 | 1.66 [-1.34, 4.66] | 1.08 | 0.278 |
|  | 400 | 1.33 [-1.30, 3.98] | 0.99 | 0.321 |
| 3000 | 200 | 3.04 [-2.35, 8.42] | 1.11 | 0.269 |
|  | 250 | 2.14 [-1.96, 6.23] | 1.02 | 0.307 |
|  | 300 | 1.49 [-1.87, 4.84] | 0.87 | 0.385 |
|  | 350 | 1.01 [-1.85, 3.86] | 0.69 | 0.488 |
|  | 400 | 0.69 [-1.81, 3.19] | 0.54 | 0.589 |

**Table S2.**
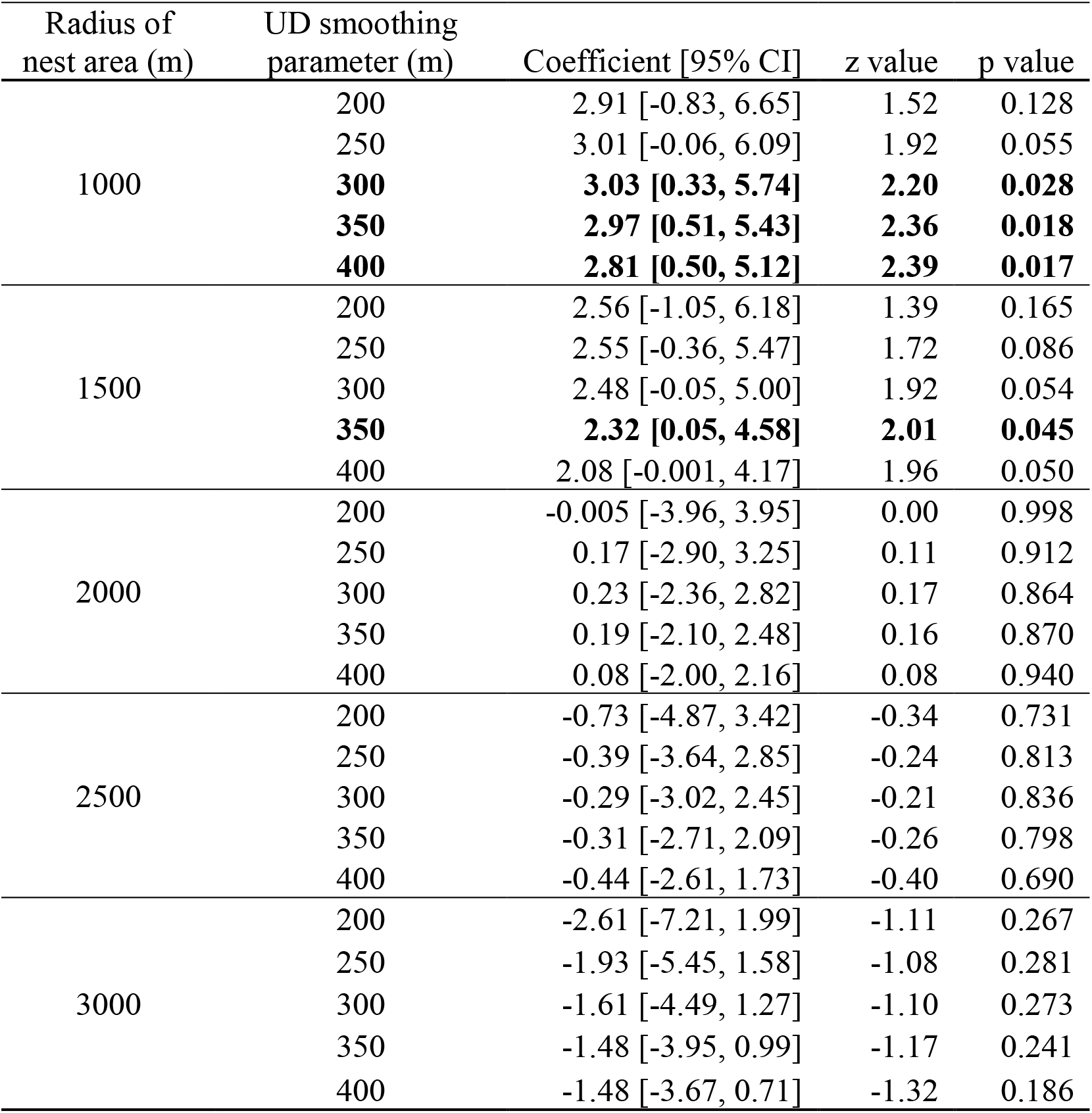
Results from conditional logistic regressions with a use-available design testing the effect of fox UD score on the nest distribution of birds that nest in micro-habitats providing partial refuge against foxes (n = 65 nests from 3 species). The fox UD score of each nest was compared to the fox UD scores of 50 random locations surrounding the nest within the nest area. Coefficient estimates are presented for 25 models, each reflecting a given size of the nest area (from 1000 to 3000 m) and UD smoothing parameter (from 200 to 400 m). Significant effects are in bold.

